# Hydrodynamic shear enables enrichment of functional tumor antigen-reactive T cells

**DOI:** 10.64898/2026.07.08.737133

**Authors:** Prem S. Subramanian, Michelle Fu, Layla C. Semaan, Asfa S. Sher, Bhupinder S. Shergill, Steven C. George, Venktesh S. Shirure

**Affiliations:** Department of Biomedical Engineering, University of California, Davis, Davis, USA; Department of Surgery, University of California, Davis, Davis, USA

**Keywords:** Tumor-reactive T cells, Hydrodynamic shear, Microfluidics, Peptide-major histocompatibility complex (pMHC), Adoptive cell therapy, Cancer immunotherapy

## Abstract

Adoptive T-cell therapies rely on the identification and expansion of rare tumor-reactive T cells, yet current enrichment strategies are limited by the low abundance of these cells and complexity of their functional enrichment. Here, we present a microfluidic platform that exploits hydrodynamic shear as a controllable parameter for enriching antigen-specific T cells through peptide-major histocompatibility complex (pMHC)-mediated capture. An eight-channel microfluidic device was engineered to simultaneously interrogate a range of wall shear stresses while maintaining uniform cell delivery, enabling systematic identification of shear conditions that maximize antigen-specific enrichment. Using engineered MART-1-specific Jurkat cells, we demonstrate that T-cell capture is jointly regulated by wall shear stress and pMHC density, with intermediate shear preferentially enriching antigen-specific cells over nonspecific binders. Translation of the optimal operating condition to a high-throughput single-shear device enabled approximately 35-fold enrichment of antigen-specific T cells from peripheral blood mononuclear cells containing only 0.05% target cells. We further show that peptide-MHC complexes isolated directly from melanoma whole-cell lysates support shear-dependent enrichment comparable to recombinant pMHCs. Finally, primary MART-1-specific CD8⁺ T cells enriched using tumor-derived pMHCs retained the ability to recognize melanoma cells and upregulated the activation marker CD137 following antigen-specific stimulation. Together, these findings establish hydrodynamic shear as an orthogonal parameter for antigen-specific T-cell enrichment and provide a framework for integrating force-based selection with tumor-derived pMHCs to isolate functional antigen-specific T cells using tumor-derived pMHCs.

## 1. Introduction

Over the past two decades, adoptive cell transfer (ACT) has provided compelling evidence that endogenous T cells can mediate durable tumor regression in patients with advanced cancer, particularly melanoma [1,2]. The recent clinical approval of tumor-infiltrating lymphocyte (TIL) therapy further underscores the therapeutic potential of naturally occurring tumor-reactive T cells in solid tumors [3,4]. However, despite these advances, response rates across solid tumors remain stubbornly modest (∼ 20%) and variable, highlighting a central limitation in current immunotherapy workflows: only a small and poorly defined fraction of T cells within heterogeneous populations both outside and within the tumor mediates effective tumor recognition and killing.

While expansion of selected T cells in a manner conducive to tumor cell killing remains an active area of research [5], perhaps the largest barrier in the field is the identification and enrichment of the rare, therapeutically relevant clonotypes that drive anti-tumor activity. Existing strategies for enriching patient-specific tumor-reactive T cells—including cytokine capture assays, tumor co-culture, and antigen-specific selection—are labor-intensive, require substantial prior knowledge of tumor antigens, or depend on large quantities of viable tumor tissue [6–8]. Even when successful, these approaches provide limited control over clonal composition and may overlook rare but potent tumoricidal T-cell populations. Consequently, there remains a need for rapid, antigen-agnostic methods to enrich functional (tumorcidal) T cells early in the manufacturing workflow, before clonal expansion.

This challenge extends across starting materials. Peripheral blood offers a minimally invasive and renewable source of T cells, enabling longitudinal sampling and broader clinical deployment. However, tumor-reactive T cells are exceedingly rare in circulation, often comprising less than 1% of peripheral blood mononuclear cells [9–11]. Conversely, while TIL products are relatively enriched for tumor-reactive cells, they remain clonally heterogeneous, and only a subset contributes meaningfully to tumor clearance [8,12]. Thus, the key limitation is not necessarily the source of lymphocytes, but the lack of robust strategies to isolate the most functionally relevant subsets from complex cellular mixtures.

Here, we introduce a fundamentally different approach to T-cell enrichment based on hydrodynamic shear or force as a controllable, biophysical determinant. Rather than selecting cells solely on the basis of equilibrium binding or phenotypic markers, this strategy enriches T cells according to their ability to sustain productive T cell receptor (TCR) interactions with peptide–major histocompatibility complex (pMHC) ligands under defined mechanical force. This concept builds on a growing body of work demonstrating that TCR signaling and antigen discrimination are intrinsically mechanosensitive processes. Mechanical forces applied across the TCR–pMHC interface can enhance specificity and signaling through mechanisms such as catch-bond formation and force-dependent kinetic proofreading [13–15].

By leveraging hydrodynamic shear within a microfluidic platform, we operationalize these principles to create a tunable selection pressure that discriminates between T cells based on functional avidity rather than simple equilibrium binding affinity. This force-based framework enables partitioning of T cells into shear-defined subsets with distinct capacities for antigen engagement and tumor cell killing. Importantly, this approach is antigen-agnostic and does not require *a priori* knowledge of tumor-specific epitopes, thereby lowering barriers to implementation across diverse tumor types and making the technology appropriate for patient-specific applications.

In this study, we test the hypothesis that calibrated hydrodynamic shear can enrich antigen-specific T cells directly from peripheral blood while preserving their functional responsiveness. Using engineered and primary MART-1-specific T-cell models together with tumor-derived peptide-MHC complexes, we establish a scalable strategy for force-based antigen-specific T-cell enrichment. Together, these findings define a new biophysical dimension for T-cell selection and provide a framework for integrating hydrodynamic enrichment with tumor-derived pMHCs for adoptive cell therapy.

## 2. Results

### 2.1 Characterization of a MART-1-specific T-cell model and validation of selective microfluidic capture

To establish a controlled model for evaluating antigen-specific T-cell capture, we generated a Jurkat-76 cell line expressing the MART-1-reactive DMF5 T-cell receptor (TCR), which recognizes the MART-1 peptide ELAGIGILTV presented by HLA-A02:01. Jurkat-76 cells lack endogenous TCR/CD3 expression, providing a clean background for expression of a defined transgenic TCR without interference from a native receptor. Following transduction and clonal selection, clone 6 (JC6) was used for all subsequent studies. Flow cytometric analysis confirmed surface expression of the transduced TCR/CD3 complex and specific binding to an *ELAGIGILTV/HLA-A02:01 pMHC tetramer* (hereafter referred to as the M1-pMHC tetramer), generated by complexing biotinylated pMHC monomers with fluorophore-conjugated streptavidin (Fig. 1A).

**Figure 1.**
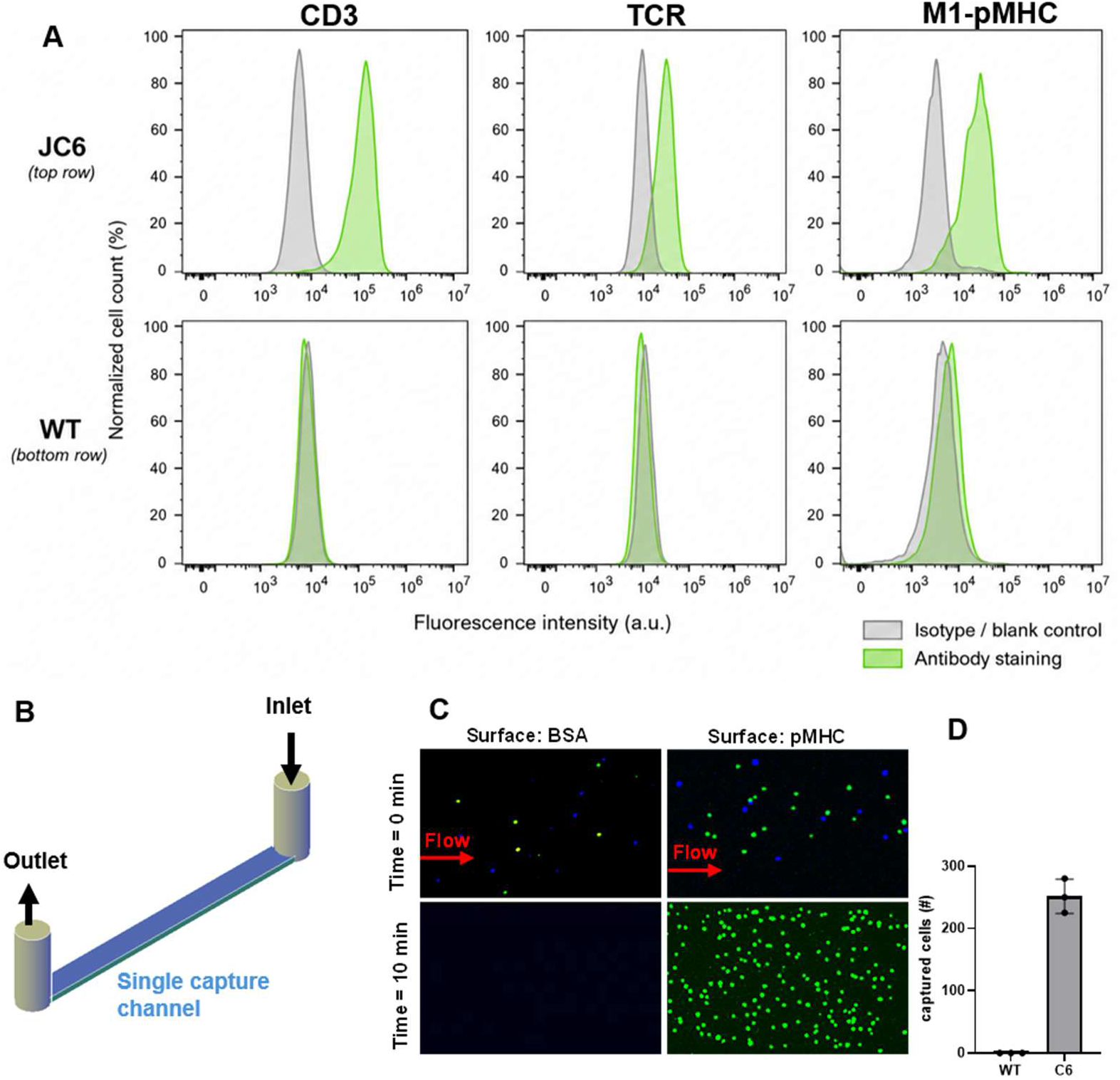
Validation of cell model and microfluidic approach for selective capture of antigen-specific T cells. (A) Flow cytometric characterization of JC6 cells (top row) and wild-type Jurkat (WT; bottom row) stained for the CD3, T-cell receptor (TCR), and MART-1/HLA-A2 tetramer (left to right). Green histograms indicate antibody or tetramer staining, and gray histograms represent the corresponding isotype or unstained controls. JC6 cells exhibit expression of the transduced MART-1-specific TCR, CD3, and MART-1 tetramer binding, whereas wild-type Jurkat cells do not bind the MART-1 tetramer. (B) Schematic of a microfluidic capture device consisting of a single functionalized capture channel connecting the inlet and outlet reservoirs. Cell suspensions are perfused through the channel under controlled flow conditions to enable selective adhesion of antigen-specific T cells. (C) Representative fluorescence microscopy images showing selective capture of JC6 cells from a mixed cell suspension during perfusion through the microfluidic channel. Green fluorescence denotes JC6 cells, whereas blue fluorescence denotes WT cells. Images were acquired before (top) and after (bottom) perfusion, illustrating preferential retention of JC6 cells on the functionalized surface while the majority of WT are removed by flow. (D) Quantification of cell capture demonstrating selective enrichment of JC6 cells over WT following perfusion through the microfluidic device. Data represent the mean ± SD.

We next established an antigen-specific capture assay using biotinylated MART-1/HLA-A*02:01 peptide-MHC (M1-pMHC) complexes containing the ELAGIGILTV peptide. The pMHCs were immobilized on NeutrAvidin-functionalized glass surfaces within a single-channel microfluidic device, and mixed suspensions of JC6 and WT Jurkat-76 cells were perfused through the channel (Fig. 1B). Representative fluorescence microscopy images demonstrated preferential retention of JC6 cells within the capture channel following perfusion, whereas WT Jurkat-76 cells were largely removed by hydrodynamic shear (Fig. 1C). Quantification of captured cells confirmed significantly greater retention of JC6 cells than WT cells (Fig. 1D), establishing that immobilized cognate pMHC complexes selectively capture antigen-specific T cells under flow.

### 2.2 Parallelized multi-shear microfluidic platform for T cell selection

To investigate the effect of hydrodynamic shear on antigen-specific T-cell capture, we designed a microfluidic platform capable of simultaneously interrogating a broad range of surface shear stresses. The platform was required to: (i) generate a controlled spectrum of wall shear stresses, and (ii) deliver cells uniformly to each capture channel so that differences in cell capture could be attributed solely to shear. To meet these design criteria, we developed an eight-channel parallel microfluidic device consisting of an inlet manifold, flow-equalizing resistors, eight capture channels, and an outlet manifold (Fig. 2A). Each capture channel had a constant height and capture area but a progressively increasing channel width, producing discrete wall shear stresses while maintaining comparable antigen presentation and available capture surface across all channels.

**Figure 2.**
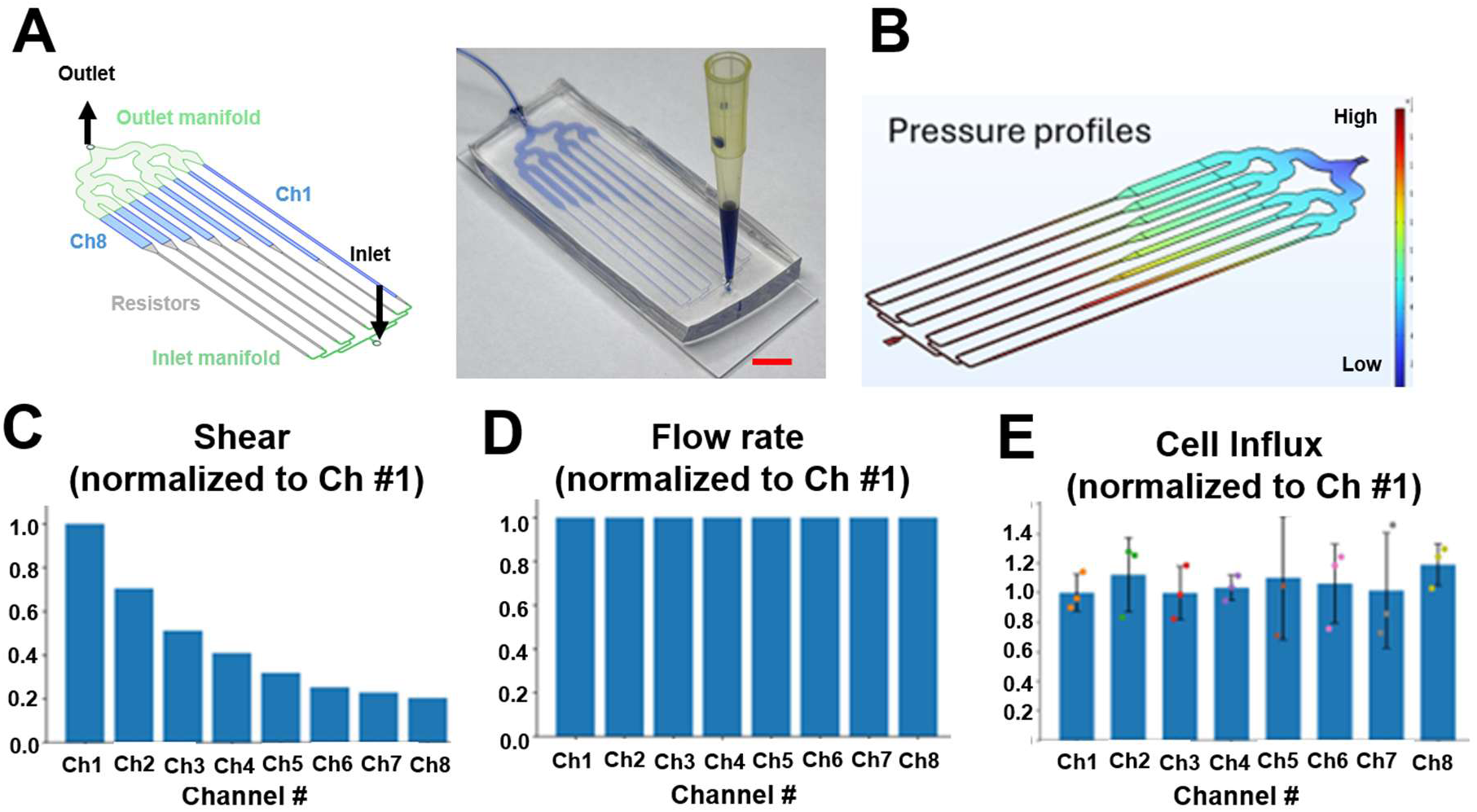
Design and validation of the parallel-channel microfluidic platform for shear-dependent T-cell interrogation. (A) Schematic and representative of the eight-channel microfluidic platform. The device consists of an inlet manifold, flow-equalizing resistors, eight parallel capture channels (Ch1-Ch8), and an outlet manifold. Capture channels have a constant height and progressively increasing widths to generate a range of wall shear stresses while maintaining the same capture area in each channel. (B) Computational fluid dynamics (CFD) simulation showing the pressure distribution throughout the microfluidic network. (C) Calculated wall shear stress in each capture channel, normalized to Channel 1, demonstrating the discrete shear profile generated across the device. (D) Simulated volumetric flow rate through each channel, normalized to Channel 1, showing uniform flow partitioning across the parallel network. (E) Experimentally measured cell influx into each capture channel, normalized to Channel 1, demonstrating uniform cell delivery despite differences in channel width and wall shear stress. Data are presented as mean ± SD.

Computational fluid dynamics (CFD) simulations confirmed the expected pressure distribution throughout the device (Fig. 2B). The design generated a discrete range of wall shear stresses across the eight channels while maintaining nearly identical volumetric flow rates in each channel (Fig. 2C,D). Experimentally, cell influx into each channel was uniform, confirming that the flow-equalizing resistors effectively partitioned cells across the parallel network despite differences in channel geometry (Fig. 2E).

Together, these results demonstrate that the platform independently modulates wall shear stress while maintaining equivalent cell delivery, enabling quantitative assessment of shear-dependent antigen-specific T-cell capture.

### 2.3 Surface M1-pMHC density and hydrodynamic shear jointly regulate antigen-specific T-cell capture

We next investigated how wall shear stress and surface M1-pMHC density influence antigen-specific T-cell capture. Sensitivity was defined as the percentage of input JC6 cells captured within the microfluidic device after perfusion. Microfluidic surfaces were functionalized with decreasing densities of M1-pMHC by diluting cognate M1-pMHC with biotin-BSA while maintaining a constant total surface coating density. The M1-pMHC densities were selected to span biologically relevant conditions, as tumor-specific pMHCs are estimated to comprise only a very small fraction of the total pMHC repertoire isolated from tumor whole-cell lysates [16,17] .

Within the low-shear regime (0.04 to 0.20 dyn cm⁻²), sensitivity depended strongly on both wall shear stress and surface M1-pMHC density (Fig. 3A). Surfaces functionalized with 100% M1-pMHC maintained a sensitivity of approximately 45 to 50% across the tested shear range. In contrast, sensitivity progressively decreased with increasing shear on surfaces coated with 1% M1-pMHC but remained above 20% at the lowest shear stresses despite a 100-fold reduction in cognate pMHC density. Sensitivity approached background levels on 0.01% M1-pMHC surfaces, and negligible sensitivity was observed on surfaces lacking M1-pMHC, confirming that cell retention resulted from specific TCR-pMHC interactions rather than nonspecific adhesion.

**Figure 3.**
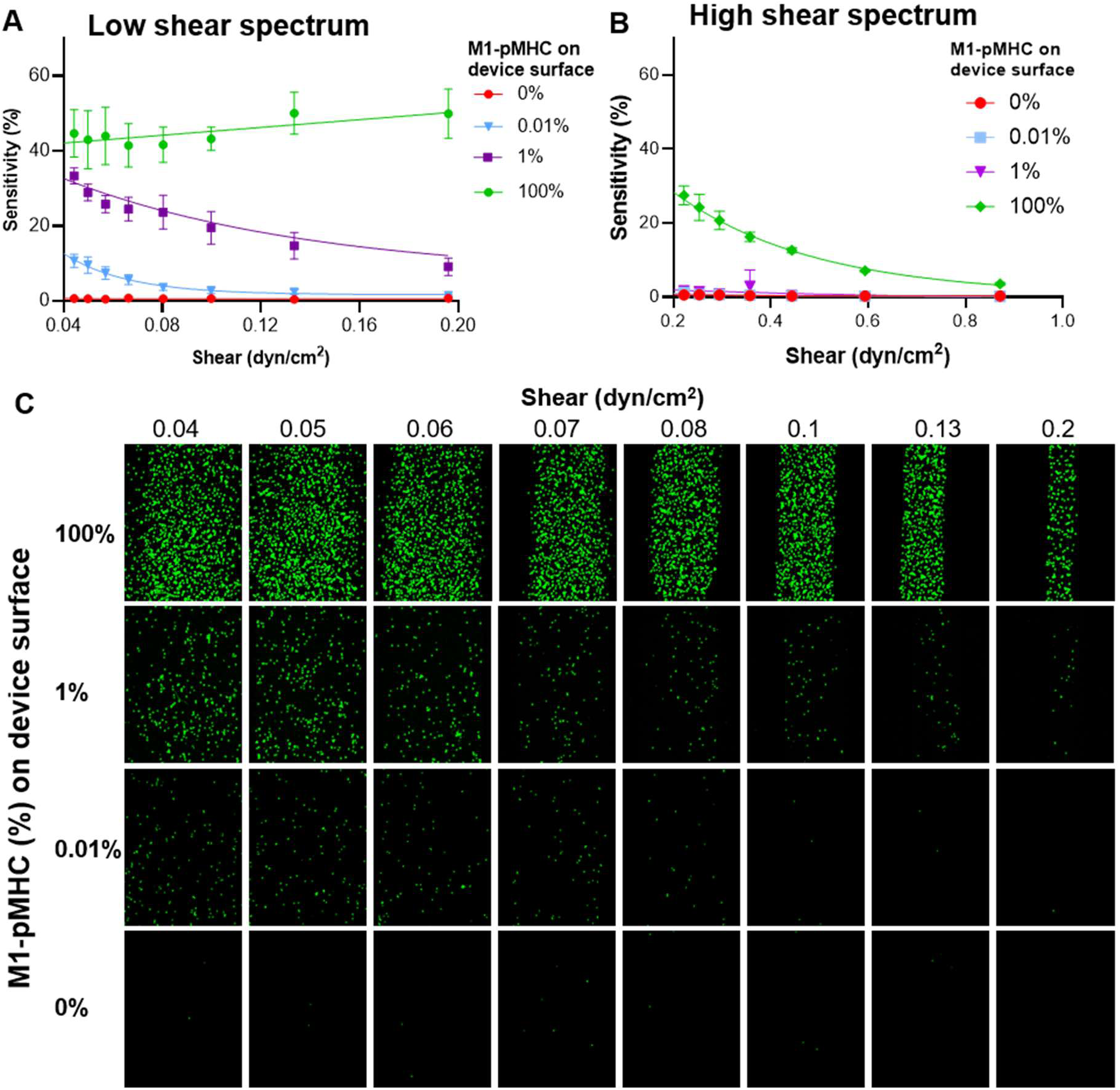
Effects of wall shear stress and surface pMHC density on antigen-specific T-cell capture. (A) Capture sensitivity of JC6 cells as a function of wall shear stress within the low-shear regime (0.04-0.20 dyn cm⁻²) for surfaces functionalized with 100%, 1%, 0.01%, or 0% MART-1/HLA-A*02:01 pMHC (M1-pMHC). Capture sensitivity decreases with increasing shear at lower pMHC densities, whereas surfaces coated with 100% pMHC maintain high capture sensitivity across the tested shear range. (B) Capture sensitivity measured over an extended high-shear range (0.20-1.0 dyn cm⁻²). High-density pMHC surfaces exhibit progressively reduced capture sensitivity with increasing shear, while negligible capture is observed on low-density and control surfaces. Data in (A) and (B) are presented as mean ± SD. (C) Representative fluorescence microscopy images of captured JC6 cells on surfaces coated with decreasing pMHC densities (rows) across increasing wall shear stresses (columns). Images qualitatively illustrate the reduction in captured cells with increasing shear and decreasing surface pMHC density. Green fluorescence indicates captured JC6 cells.

To determine the effect of higher hydrodynamic forces on antigen-specific capture, we extended the analysis to wall shear stresses ranging from 0.20 to 1.0 dyn cm⁻² (Fig. 3B). Even on surfaces coated with 100% M1-pMHC, sensitivity decreased monotonically with increasing shear and approached background levels at the highest shear stresses tested. As expected, 1%, 0.01%, and control surfaces exhibited negligible sensitivity throughout this higher shear regime. Representative fluorescence microscopy images qualitatively illustrate the progressive reduction in captured JC6 cells with increasing wall shear stress and decreasing surface M1-pMHC density (Fig. 3C).

Collectively, these findings demonstrate that antigen-specific T-cell sensitivity is jointly regulated by surface M1-pMHC density and hydrodynamic shear. Importantly, M1-pMHC densities approximating those expected for tumor-specific pMHCs in whole-cell lysate preparations (approximately 1%) still supported greater than 20% sensitivity under low-shear conditions. These findings suggest that tumor-derived pMHC preparations should retain sufficient antigen density to enable efficient capture of cognate T cells and provide the rationale for subsequent studies using tumor-derived pMHCs.

### 2.4 Hydrodynamic shear enables enrichment of antigen-specific T cells from complex cell mixtures

Having established the effects of M1-pMHC density on antigen-specific capture, we next investigated whether hydrodynamic shear could be exploited to enrich rare antigen-specific T cells from a heterogeneous cell population. JC6 cells were spiked into human PBMCs at an input frequency of approximately 0.5%, representing the low frequency of tumor-reactive T cells typically found in peripheral blood, and perfused over surfaces presenting 1% M1-pMHC diluted within a background of nonspecific pMHCs to model the low abundance of tumor-specific pMHCs expected in tumor-derived whole-cell lysates (Fig. 4A).

**Figure 4.**
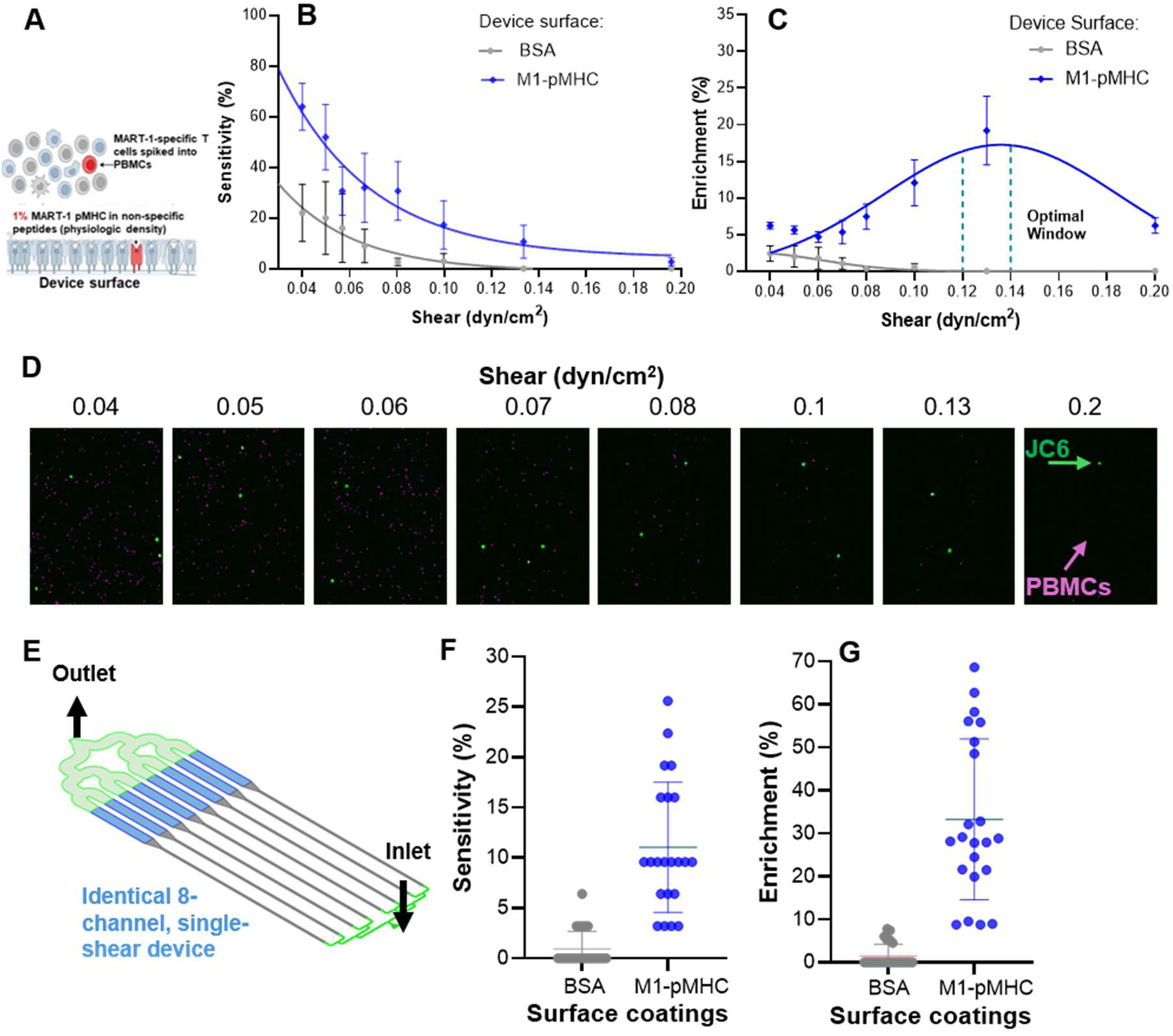
Hydrodynamic shear enables enrichment of antigen-specific T cells from a PBMC background. (A) Schematic of the experimental model in which JC6 cells were spiked into human peripheral blood mononuclear cells (PBMCs) and perfused over surfaces functionalized with 1% M1-pMHC diluted in nonspecific pMHCs to mimic the low abundance of tumor-specific pMHCs expected in tumor-derived whole-cell lysates. (B) Sensitivity of JC6 cells (blue) and PBMCs (gray) as a function of wall shear stress. Sensitivity was calculated as the percentage of input cells captured within the microfluidic device after perfusion. (C) Fold enrichment of JC6 cells relative to the input population as a function of wall shear stress. Maximum enrichment was observed at an intermediate wall shear stress (∼0.13 dyn cm⁻²), which was selected for subsequent validation in the single-shear device. Data in (B) and (C) are presented as mean ± SD. (D) Representative fluorescence microscopy images of captured cells across increasing wall shear stresses. JC6 cells are shown in green and PBMCs in magenta. Increasing wall shear progressively removes nonspecifically captured PBMCs while retaining antigen-specific JC6 cells over an intermediate shear range. (E) Schematic of the single-shear microfluidic device consisting of eight identical capture channels operated at the selected wall shear stress of 0.13 dyn cm⁻², identified from the shear optimization experiments in (C). (F) Sensitivity of JC6 cells captured using the single-shear microfluidic device operated at 0.13 dyn cm⁻². (G) Fold enrichment of JC6 cells captured using the single-shear microfluidic device operated at 0.13 dyn cm⁻². The error bars indicate mean ± SD.

Sensitivity and fold enrichment were evaluated as a function of wall shear stress. Sensitivity was defined as the percentage of input JC6 cells captured within the microfluidic device after perfusion. Fold enrichment was defined as the fraction of JC6 cells among captured cells divided by the fraction of JC6 cells in the input perfusate. As expected, JC6 sensitivity decreased with increasing wall shear stress (Fig. 4B). In contrast, negligible capture was observed on Biotin-BSA control surfaces across the entire shear range, confirming that cell capture was mediated by specific M1-pMHC interactions rather than nonspecific surface adhesion.

Despite the gradual reduction in JC6 sensitivity, fold enrichment exhibited a nonmonotonic dependence on wall shear stress (Fig. 4C). Enrichment increased with increasing shear, reached a maximum at an intermediate wall shear stress of approximately 0.13 dyn cm⁻², and declined at higher shear stresses as increasing numbers of antigen-specific JC6 cells were detached. Representative fluorescence microscopy images qualitatively illustrate this transition, showing progressive removal of PBMCs while retaining antigen-specific JC6 cells within the intermediate shear regime (Fig. 4D).

The wall shear stress corresponding to maximum enrichment (0.13 dyn cm⁻²) was subsequently implemented in a high-throughput single-shear microfluidic device containing eight identical capture channels (Fig. 4E). Unlike the shear-screening device, this format enabled all captured cells to be recovered under a single optimized operating condition, substantially increasing throughput and facilitating enrichment experiments with rarer cell populations. Using this device, JC6 cells spiked into PBMCs at an input frequency of 0.05% were captured with robust sensitivity while maintaining substantial fold enrichment (Fig. 4F,G). Together, these results demonstrate that hydrodynamic shear can be tuned to selectively enrich rare antigen-specific T cells from complex cellular backgrounds and that the identified operating condition can be readily translated to a scalable, higher-throughput microfluidic platform.

### 2.5 Tumor-derived pMHCs support hydrodynamic enrichment of antigen-specific T cells

Having established the shear-dependent enrichment strategy using recombinant M1-pMHC, we next investigated whether tumor-derived pMHCs could similarly support antigen-specific T-cell capture. As a source of naturally processed tumor antigens, we used the HLA-A*02:01-positive melanoma cell line MEL624, which expresses the MART-1 antigen. Intracellular flow cytometry confirmed robust MART-1 expression in MEL624 cells (Fig. 5A). pMHC complexes were isolated from MEL624 whole-cell lysates by immunoaffinity purification, biotinylated, and immobilized onto NeutrAvidin-functionalized microfluidic surfaces (Fig. 5B).

**Figure 5.**
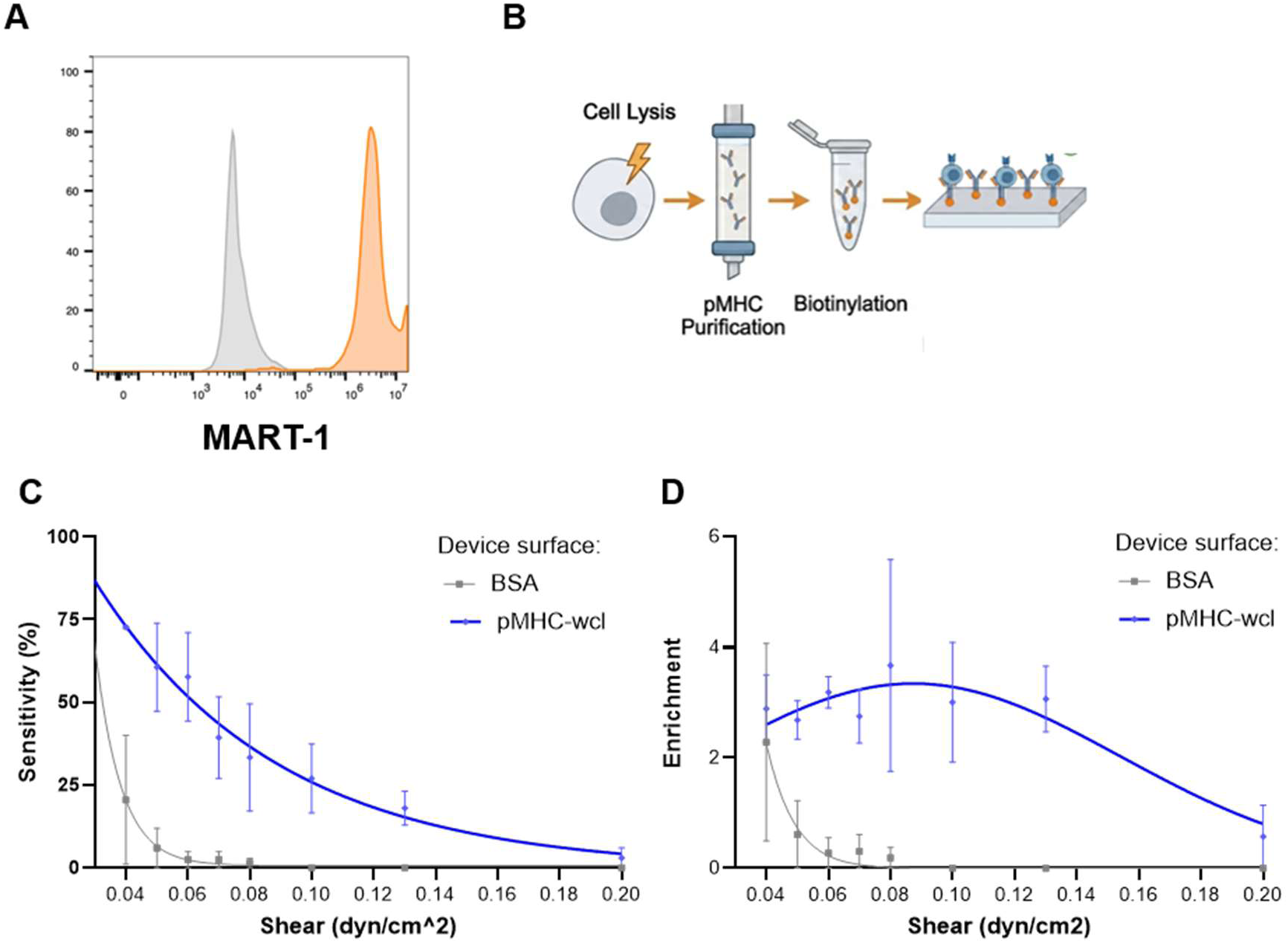
Tumor-derived pMHCs enable antigen-specific T-cell capture and enrichment under hydrodynamic shear. (A) Flow cytometric validation of intracellular MART-1 expression in MEL624 melanoma cells. Orange histogram represents MART-1 staining and gray histogram represents the corresponding isotype control. (B) Schematic illustrating the isolation of peptide-MHC (pMHC) complexes from MEL624 whole-cell lysates by immunoaffinity purification, followed by biotinylation and immobilization on NeutrAvidin-functionalized microfluidic surfaces. (C,D) JC6 cells were spiked into PBMCs at an input frequency of 0.5% and perfused over microfluidic surfaces functionalized with tumor-derived pMHCs isolated from MEL624 melanoma cells (blue) or biotin-BSA control surfaces (gray). Sensitivity (C) and fold enrichment (D) are shown as a function of wall shear stress. Maximum fold enrichment was observed at an intermediate wall shear stress. Data in (C) and (D) are presented as mean ± SD.

JC6 cells were spiked into PBMCs at an input frequency of approximately 0.5% and perfused over tumor-derived pMHC-functionalized surfaces. Sensitivity and fold enrichment were evaluated as a function of wall shear stress, with Biotin-BSA-coated surfaces serving as negative controls (Fig. 5C,D). Consistent with results obtained using recombinant M1-pMHC, JC6 sensitivity decreased progressively with increasing wall shear stress, whereas negligible capture was observed on Biotin-BSA control surfaces across the entire shear range, confirming that cell capture required tumor-derived pMHCs (Fig. 5C).

Fold enrichment again exhibited a nonmonotonic dependence on wall shear stress, reaching a maximum at an intermediate shear before declining at higher shear stresses as antigen-specific JC6 cells were increasingly detached (Fig. 5D). Although the maximum fold enrichment achieved with tumor-derived pMHCs was lower than that observed using recombinant M1-pMHC (Fig. 4C), the characteristic shear-dependent enrichment profile was preserved. These findings demonstrate that naturally processed tumor-derived pMHCs retain sufficient antigenic information to selectively capture and enrich cognate T cells under hydrodynamic shear, providing the foundation for extending this approach to patient-derived tumor samples.

### 2.6 Tumor-derived pMHCs enrich functional primary antigen-specific T cells

To determine whether tumor-derived pMHCs could enrich functional primary antigen-specific T cells, commercially available MART-1-specific CD8⁺ T cells were spiked into HLA-A*02:01-matched donor PBMCs at an input frequency of approximately 0.5% and perfused over microfluidic surfaces functionalized with either recombinant M1-pMHC or MEL624 whole-cell lysate (WCL)-derived pMHCs (Fig. 6A). The input population and cells recovered following microfluidic capture were subsequently co-cultured with MEL624 melanoma cells for 24 h before analysis of antigen specificity and activation by flow cytometry.

**Figure 6.**
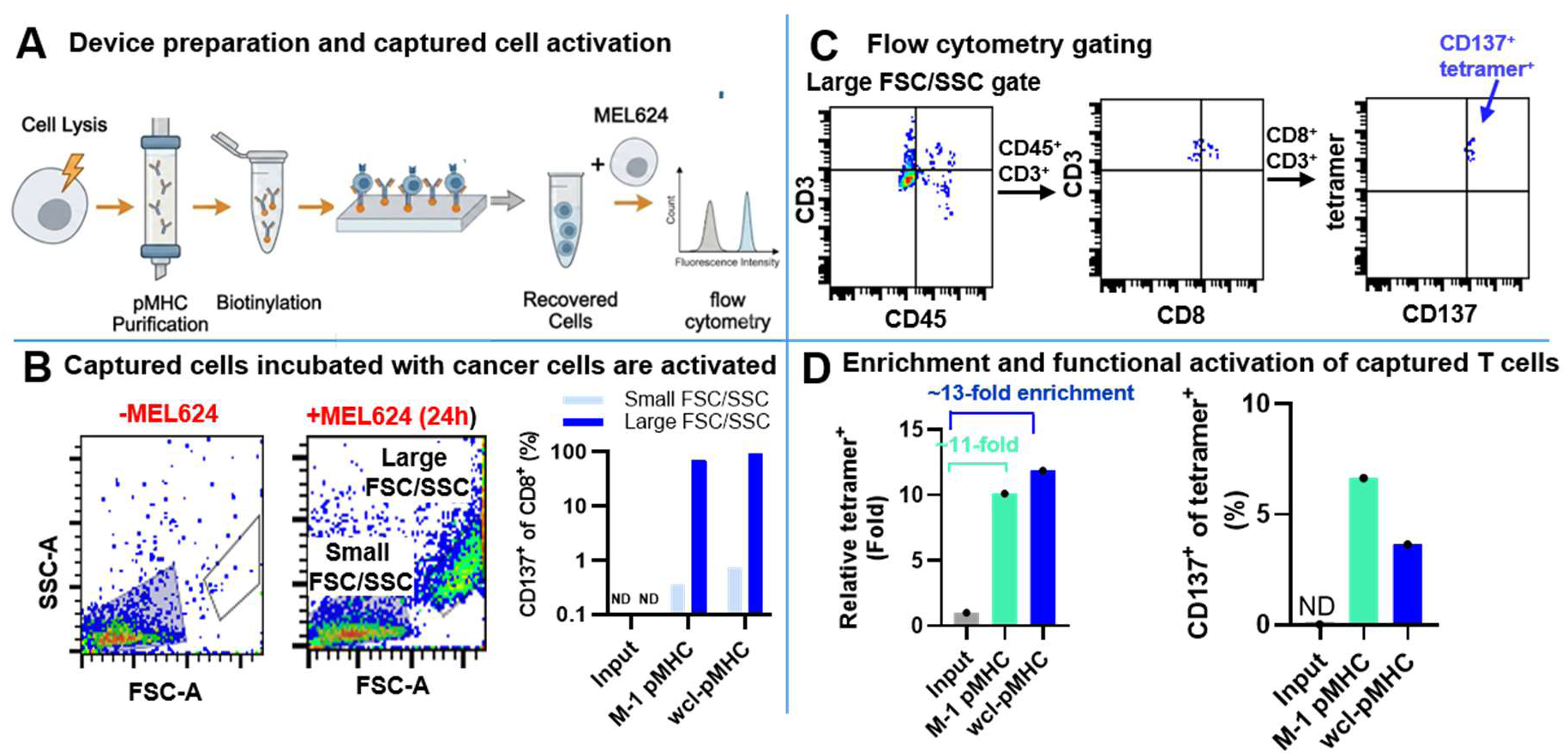
Tumor-derived pMHCs enrich functional primary antigen-specific T cells. (A) Experimental workflow. Peptide-MHC (pMHC) complexes were isolated from MEL624 melanoma whole-cell lysates, biotinylated, and immobilized on NeutrAvidin-functionalized microfluidic surfaces. Commercially available primary MART-1-specific CD8⁺ T cells were spiked into HLA-A*02:01-matched donor PBMCs at an input frequency of 0.5%, captured on tumor-derived pMHC surfaces, and recovered cells were co-cultured with MEL624 melanoma cells for 24 h prior to flow cytometric analysis. (B) Representative forward scatter (FSC) and side scatter (SSC) plots before (left) and after (right) 24 h co-culture with MEL624 cells. Co-culture resulted in the appearance of a distinct large FSC/SSC population in addition to the lymphocyte-sized population. Quantification of M-1 pMHC tetramer-positive cells within the small and large FSC/SSC populations before and after co-culture is shown at right. (C) Flow cytometry gating strategy used to identify M-1 pMHC tetramer-positive T cells and quantify CD137 expression. Representative M-1 pMHC tetramer and CD137 dot plots are shown for the input population, cells recovered from the microfluidic device, and recovered cells following 24 h co-culture with MEL624 cells. (D) Quantification of M1-pMHC tetramer-positive T-cell enrichment relative to the input population following microfluidic capture on surfaces functionalized with either recombinant M1-pMHC or MEL624 whole-cell lysate (WCL)-derived pMHCs. The input population and both captured populations were subsequently co-cultured with MEL624 melanoma cells for 24 h, after which M1-pMHC tetramer-positive T cells (left) and CD137 expression within the M1-pMHC tetramer-positive population (right) were quantified by flow cytometry.

Antigen-specific T cells and their activation status were analyzed using the flow cytometry gating strategy shown in Fig. 6C. Following co-culture, a distinct population of larger FSC/SSC cells emerged in addition to the lymphocyte-sized population (Fig. 6B). This large FSC/SSC population was enriched for CD137-positive cells recovered from both microfluidic capture surfaces, whereas CD137-positive cells were largely absent from the input population following the same co-culture conditions.

Microfluidic capture on both recombinant M1-pMHC and WCL-derived pMHC surfaces substantially increased the frequency of M1-pMHC tetramer-positive T cells relative to the input population, resulting in approximately 11-fold and 13-fold enrichment, respectively (Fig. 6D). Importantly, tetramer-positive T cells recovered from both capture surfaces remained functionally responsive, as demonstrated by robust CD137 upregulation following co-culture with MEL624 cells (Fig. 6D). These findings demonstrate that tumor-derived pMHC-functionalized microfluidic surfaces not only enrich primary antigen-specific T cells from complex PBMC populations but also preserve their ability to recognize naturally processed tumor antigens and become activated.

## 3. Discussion

In this study, we introduce a hydrodynamic force-based strategy for enriching antigen-specific T cells using peptide-MHC (pMHC)-functionalized microfluidic surfaces. Rather than relying solely on affinity- or phenotype-based selection, our approach exploits the force dependence of TCR-pMHC interactions to selectively retain antigen-specific T cells under defined hydrodynamic conditions. Using engineered model systems, recombinant peptide-MHC complexes, tumor-derived pMHCs, and primary human MART-1-specific T cells, we demonstrate that intermediate wall shear stresses maximize enrichment while maintaining antigen-specific capture. Importantly, T cells recovered using tumor-derived pMHCs remained functionally responsive to melanoma cells, indicating that the enrichment process preserves their ability to recognize naturally processed tumor antigens.

A key implication of this work is the potential to address a longstanding bottleneck in adoptive cell therapy: the identification and enrichment of rare tumor-reactive T cells from highly heterogeneous starting populations. Using the multi-shear platform, we identified a hydrodynamic operating window that maximized antigen-specific enrichment while maintaining cell recovery. Importantly, this operating condition was successfully translated to a high-throughput single-shear microfluidic device, which achieved approximately 35-fold enrichment when antigen-specific T cells were present at only 0.05% of the input population. This result demonstrates that the mechanically defined operating window remains effective under biologically relevant rare-cell conditions and highlights the scalability of the approach. Such scalability is particularly relevant for peripheral blood-based immunotherapies, where tumor-reactive T cells are exceedingly rare but represent an attractive, minimally invasive source for adoptive cell therapy.

The observed dependence of enrichment on wall shear stress is consistent with the force-sensitive nature of TCR-pMHC interactions. At low shear stresses, both specific and weakly adherent cells remain associated with the capture surface, resulting in limited enrichment. Increasing wall shear preferentially removes weakly bound or nonspecifically adherent cells while retaining antigen-specific T cells, thereby increasing enrichment. At higher shear stresses, however, hydrodynamic forces exceed the strength of productive TCR-pMHC interactions, leading to loss of antigen-specific cells and reduced enrichment. This balance between retention and detachment defines a mechanical operating window for antigen-specific T-cell enrichment.

Despite these encouraging results, several considerations will influence the translation of this approach to clinical samples. The composition and abundance of tumor-derived pMHCs are expected to vary substantially across patients, tumor types, and even individual lesions [16–18]. While MEL624-derived pMHCs demonstrated that naturally processed tumor antigens can support shear-dependent enrichment, patient tumors are likely to exhibit greater heterogeneity in antigen repertoire, MHC expression, and peptide presentation [16,17]. Consequently, the optimal hydrodynamic operating window may differ between samples and require calibration for individual tumors. One potential strategy to improve enrichment is to increase the fraction of tumor-derived pMHCs before surface functionalization. For example, enrichment of tumor cells from surgical specimens prior to pMHC isolation could reduce contamination from stromal and immune cells, thereby increasing the relative abundance of tumor-associated pMHCs and improving antigen-specific capture.

A second consideration is the diversity of circulating tumor-reactive T cells. Unlike the engineered and primary MART-1-specific T-cell models used here, endogenous tumor-reactive T cells comprise heterogeneous populations that differ in TCR affinity, activation state, differentiation status, and antigen specificity. These biological differences may influence the mechanical stability of TCR-pMHC interactions and, consequently, the efficiency of shear-based enrichment. Although the present study establishes proof-of-concept using defined antigen-specific T-cell populations, extending this approach to matched patient tumor and peripheral blood samples will be important to evaluate its performance in clinically relevant settings.

Although hydrodynamic shear substantially reduced nonspecific cell retention, it is unlikely to completely eliminate background capture. Weak TCR-pMHC interactions, cross-reactive T cells, or adhesion mediated through non-TCR receptors may contribute to residual nonspecific capture, particularly when using complex tumor-derived pMHC repertoires. This challenge is analogous to current tumor-infiltrating lymphocyte (TIL) therapies, in which only a fraction of the expanded product is thought to be truly tumor reactive, while the remaining cells may represent bystander or unrelated T-cell populations [8,19]. By enriching antigen-specific T cells prior to expansion, hydrodynamic selection has the potential to increase the fraction of antigen-specific T cells within the final therapeutic product. Additional optimization of surface chemistry, pMHC presentation, and downstream selection strategies may further improve specificity.

Finally, enrichment alone does not ensure successful manufacture of therapeutic cell products. The ability of captured T cells to expand, maintain functionality, and persist following ex vivo culture remains an important consideration for adoptive cell therapy [20–22]. Our finding that enriched primary antigen-specific T cells retained the capacity to recognize naturally processed tumor antigens and upregulate CD137 following tumor stimulation demonstrates that hydrodynamic enrichment is compatible with downstream functional responses. Future studies integrating this approach with ex vivo expansion, TCR repertoire sequencing, and single-cell transcriptomic profiling will provide further insight into the phenotype, clonal diversity, and therapeutic potential of mechanically enriched T-cell populations.

## 4. Conclusion

In conclusion, this work establishes hydrodynamic shear as a controllable parameter for enriching antigen-specific T cells using both recombinant and tumor-derived pMHCs. The ability to recover primary T cells that remain responsive to naturally processed tumor antigens demonstrates the feasibility of coupling force-based selection with functional immunotherapy workflows. Future studies using matched patient tumors and peripheral blood will determine the extent to which this strategy can enrich naturally occurring tumor-reactive T-cell repertoires for adoptive cell therapy.

## 5. Methods

### 5.1 Microfabrication and Surface Device Preparation

Microfluidic device fabrication utilized standard methods of soft lithography. Microfluidic channel geometry was fabricated on a silicon wafer master mold using SU-8 photolithography. Polydimethylsioxane (PDMS; Sigma-Aldrich, Cat: 504823) was prepared by combining Sylgard 184 elastomer base and curing agent at a 10:1 ratio and evenly mixed. After pouring the resultant mixture, the device was cured at 60 °C for at least 4 hours. Glass slides (Thermo Fisher Scientific) were rinsed thoroughly in purified water and ethanol. The cured PDMS containing the channel features was then peeled from the mold and plasma bonded to the glass slide. The resultant devices were cured overnight at 120 C to seal and strengthen the covalent bond between the PDMS device and the glass slide.

Microfluidic devices were chemically functionalized to enable immobilization of peptide-major histocompatibility complex (pMHC) molecules for T-cell capture experiments. Prior to functionalization, devices were plasma treated to activate the channel surfaces. The channels were then sequentially treated with 3.65% (v/v) silane for 1 h, 1 mM GMBS (N-γ-maleimidobutyryloxy succinimide ester) for 30 min, and NeutrAvidin for 1 h.

Biotinylated MART-1 peptide-MHC was obtained from the NIH Tetramer Core Facility (GA). Biotinylated bovine serum albumin (biotin-BSA) was prepared separately. The two solutions were mixed at defined ratios to achieve the desired surface density of pMHC.

### 5.2 Generation of a MART-1-specific Jurkat-76 subclone

Jurkat 76 cells were grown at 2*10^5^ cells in 2 ml media containing 4 μg/mL l polybrene and 100 ml lentivirus titer (**Supplemental Fig. 1**) were spun in 15 ml conical tunes at 800g for 1h at 32 C. The pellet was resuspended in 2 ml media and grown for 3 days in 6wp. 2 ml fresh media with 4 μg/mL Puromycin was added to each well. Cells were grown for 3 more days. On day 6 post transduction - subclone 3X96wp (calculated 5 cells in 100 ml in each well).

Cells were grown for 3 days without antibiotics, then 3 more days with Puromycin (2 mg/ml). Cells in each well then sorted by FACS to select the brightest mCherry population. Subclones were then grown for 6 more days in Puromycin (2 mg/ml).

Twenty-four subclones were screened by flow cytometry for mCherry, TCRαβ, and CD3 expression. Clone 6 (JC6), which exhibited the highest and most uniform expression, was selected for all subsequent experiments.

### 5.3 Cell Staining

PBMCs were isolated from healthy volunteers as previously described. Cryopreserved PBMCs were rapidly thawed, diluted in pre-warmed RPMI 1640 medium (Gibco, Thermo Fisher Scientific), washed by centrifugation (400 × g, 5 min) to remove DMSO, and resuspended in DPBS (-/-). Cell concentration and viability were determined by trypan blue exclusion using an automated cell counter (Countess, Thermo Fisher Scientific).

PBMCs were labeled with CellTracker Deep Red (Invitrogen, Thermo Fisher Scientific; Cat. No. C34565) according to the manufacturer’s instructions. Briefly, cells were incubated with dye for 30-60 min at 37 °C, washed twice with DPBS, and staining was confirmed by fluorescence microscopy prior to experimentation. Jurkat C6 cells were labeled using the same protocol with CellTracker Green (Invitrogen, Thermo Fisher Scientific; Cat. No. C7025).

### 5.4 Cell Perfusion Through the Microfluidic Device

Prior to each experiment, microfluidic devices were equilibrated to room temperature, washed with DPBS (-/-) for 10 min, and connected to a syringe pump (Harvard Apparatus) operating in withdrawal mode. Devices were mounted on an FV1200 Fluoview confocal laser scanning microscope (Olympus) for real-time imaging during perfusion. Cells were suspended at 5 × 10⁵ cells/mL and perfused through the devices at the specified flow rates. Following perfusion, devices were washed with DPBS for 30 min at the same flow rate to remove non-adherent cells before imaging.

For downstream analyses, adherent cells were recovered by gentle pipetting through the outlet, maintained in warm RPMI medium, and processed for flow cytometry as described below.

### 5.5 Flow Cytometry

For Jurkat C6 and wild-type (WT) cells, approximately 2 × 10⁵ cells per sample were processed using the same staining protocol. Unstained samples were included as controls for flow cytometry.

For intracellular MART-1 detection, cells were fixed, permeabilized, and stained using the Cyto-Fast™ Fix/Perm Kit according to the manufacturer’s instructions. The following antibodies were evaluated: anti-MART-1 TCR-like antibody (Creative Biolabs), anti-Melan-A clone A103 (Santa Cruz Biotechnology), and mouse anti-human MART-1/Melan-A Alexa Fluor 647.

For multicolor flow cytometry experiments, compensation was performed using single-stained compensation beads and fluorescence minus one (FMO) controls were included to establish gating for low-frequency cell populations.

### 5.6 Melanoma Cell Culture and Whole cell protein isolation

MEL624 melanoma cells, which express MART-1, were cultured in DMEM supplemented with fetal bovine serum and penicillin-streptomycin until confluent. Cells were detached using Versene (DPBS -/- containing 2 mM EDTA), collected, and lysed in 1% (w/v) digitonin lysis buffer containing Tris-HCl, EDTA, NaCl, and Halt protease inhibitor. Lysates were incubated on ice for 40 min, clarified by centrifugation (16,000-20,000 × g, 20 min), and the resulting whole-cell lysates were collected. Total protein concentration was determined using the Micro BCA Protein Assay Kit (Thermo Fisher Scientific), and lysates were stored at −80 °C until use.

### 5.7 Isolation of pMHCs and Surface Coating

pMHC complexes were isolated from MEL624 whole-cell lysates using the Pierce Co-Immunoprecipitation Kit. To preserve peptide-MHC complexes, the Gentle Ag/Ab Elution Buffer (pH 6.6) was used for elution, after which eluates were neutralized to pH 7.2-7.3 with 1 M NaOH. Purified pMHCs were biotinylated using the EZ-Link Sulfo-NHS-Biotin Labeling Kit, and excess biotin was removed using Zeba Spin Desalting Columns. All procedures were performed on ice to minimize peptide dissociation. Biotinylated pMHCs were stored at 4 °C until use.

Microfluidic devices were coated with biotinylated pMHCs by incubating the pMHC solution within NeutrAvidin-functionalized channels at 4 °C with periodic recirculation to promote uniform surface coverage. Coated devices were stored at 4 °C and used within 2 days.

### 5.8 Statistical Analysis

Data are presented as means ± standard deviation. Statistical analysis was performed using one-way ANOVA with post-hoc Tukey test, two-way ANOVA with Tukey’s multiple comparisons test, or unpaired t-test when necessary.

## Acknowledgements

This work was supported by a grant from the National Institutes of Health (R61 CA278531 (VSS). We would also like to acknowledge support from the Provost Undergraduate Fellowship which supported summer research on this project for LS and and technical assistance from Dr. Sergey Yechikov for the creation of the mCherry DMF5 MART-1 plasmid and stable subclone (C6) of the Jurkat 76 cell line. We thank the NIH Tetramer Core Facility for providing the MART-1 tetramers. We thank Center for Nano Micro Manufacturing facility at UC Davis for assistance with microfabrication.

## Supplementary Material

**Supplementary Figure 1.**
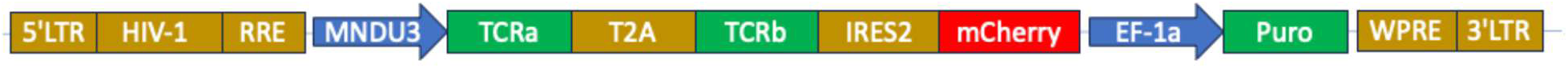
Plasmid **Map and Sequence**

## Notes

### Competing Interest Statement

The authors have declared no competing interest.

