## Supplementary figures and images for "Hydrodynamic shear enables enrichment of functional tumor antigen-reactive T cells"

### Supplemental Fig. 1

#### Supplementary Material

**Supplementary Figure 1.** Plasmid Map and Sequence

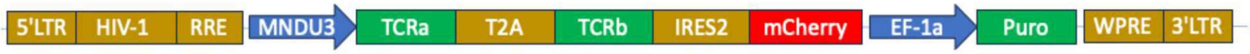
